# Food web interaction strength distributions are conserved by greater variation between than within predator-prey pairs

**DOI:** 10.1101/461921

**Authors:** Daniel L. Preston, Landon P. Falke, Jeremy S. Henderson, Mark Novak

## Abstract

Species interactions in food webs are usually recognized as dynamic, varying across species, space and time due to biotic and abiotic drivers. Yet food webs also show emergent properties that appear consistent, such as a skewed frequency distribution of interaction strengths (many weak, few strong). Reconciling these two properties requires an understanding of the variation in pairwise interaction strengths and its underlying mechanisms. We estimated stream sculpin feeding rates in three seasons at nine sites in Oregon to examine variation in trophic interaction strengths both across and within predator-prey pairs. We considered predator and prey densities, prey body mass, and abiotic factors as putative drivers of within-pair variation over space and time. We hypothesized that consistently skewed interaction strength distributions could result if individual interaction strengths show relatively little variation, or alternatively, if interaction strengths vary but shift in ways that conserve their overall frequency distribution. We show that feeding rate distributions remained consistently and positively skewed across all sites and seasons. The mean coefficient of variation in feeding rates within each of 25 focal species pairs across surveys was less than half the mean coefficient of variation seen across species pairs within a given survey. The rank order of feeding rates also remained relatively conserved across streams, seasons and individual surveys. On average, feeding rates on each prey taxon nonetheless varied by a hundredfold across surveys, with some feeding rates showing more variation in space and others in time. For most species pairs, feeding rates increased with prey density and decreased with high stream flows and low water temperatures. For nearly half of all species pairs, factors other than prey density explained the most variation, indicating that the strength of density dependence in feeding rates can vary greatly among a generalist predator’s prey species. Our findings show that although individual interaction strengths exhibit considerable variation in space and time, they can nonetheless remain relatively consistent, and thus predictable, compared to the even larger variation that occurs across species pairs. These insights help reconcile how the skewed nature of interaction strength distributions can persist in highly dynamic food webs.

## Introduction

Most attributes of food webs – including species composition and abundances, network topology, and interaction strengths – vary in space and time (Menge et al. 1994, Polis et al. 1996). Deterministic drivers of food web variation include both biotic factors such as species introductions or extirpations, population cycles, and organism life history traits (e.g., Boutin et al. 1995, Vander Zanden et al. 1999, de Roos et al. 2003), and abiotic factors such as temperature, nutrients, hydrology, light, and substrate (e.g., Menge 2000, Power et al. 2008, Byers et al. 2017). For example, migrations of anadromous fish can drive predictable seasonal changes in web topology and energy flow (Naiman et al. 2002); tropical storms can rapidly alter interaction strengths on islands (Spiller and Schoener 2007); and climate change is leading to wholesale food web alterations on global scales (Woodward et al. 2010). While increasingly recognized, spatial and temporal food web variation present challenges to predicting and managing community dynamics, particularly in species-rich communities where the relevant intrinsic and extrinsic drivers are poorly resolved (Tylianakis et al. 2008).

Although a large body of research shows that food webs are inherently variable, some empirical food web patterns appear to be relatively consistent in space and time (Mora et al. 2018). Among these is the skewed frequency distribution of interaction strengths (few strong and many weak) that has been documented in nearly all studies with field-based quantitative interaction strength measures (e.g., Paine 1992, Fagan and Hurd 1994, de Ruiter et al. 1995, Raffaelli and Hall 1996, Wootton 1997, Woodward et al. 2005, Schleuning et al. 2011, Cross et al. 2013, Bellmore et al. 2015). This pattern appears insensitive to ecosystem type, network complexity, the measure of interaction strength used, and even the type of interaction under study (Wootton and Emmerson 2005, Vázquez et al. 2012). The persistence of the skewed distribution of interaction strengths suggests that either: 1) despite being variable, the strength of each pairwise species interaction shows consistency relative to the variation seen across co-occurring interactions, or 2) the relative position of each pairwise interaction along the distribution is dynamic, but with a distribution-conserving fraction of interactions shifting from strong to weak and vice versa. The latter might occur, for example, if predators are limited by a maximum total feeding rate across all of their prey. Most quantitative measures of species interaction strength lack the spatial or temporal replication to test these ideas for multiple co-occurring interactions in nature.

Estimates of predator feeding rates are useful for addressing the extent to which species interaction strengths and their frequency distributions are dynamic or consistent over space and time. Moreover, there is a rich literature that seeks to mechanistically describe the factors driving variation in feeding rates (Jeschke et al. 2002). For example, functional response models generally predict that feeding rates should increase (often nonlinearly) with prey density (Holling 1959), such that fluctuations in prey should be a primary factor determining variation. Predator density, predator and prey traits (e.g., body size), and environmental conditions are also linked to variation in feeding rates (e.g., Skalski and Gilliam 2001, Rall et al. 2012, Kalinoski and DeLong 2016), with changes in each having the potential to alter interaction strengths and their frequency distribution in space and time.

In the present study we address two related questions using replicated *in situ* feeding rate estimates of a focal generalist predator, the reticulate sculpin (*Cottus perplexus*). First, we ask how dynamic are prey-specific sculpin feeding rates in space and time? To address this question, we use the variation seen in sculpin feeding rates across their diverse prey community as a relative measure to compare against the variation seen within species pairs over space and time. Variation within species pairs across space and time that is consistently less than variation across species pairs at a given point in space and time would suggest that pairwise species interaction strengths show consistency, which could underlie the consistency of community-wide interaction strength frequency distributions. A conserved rank order of prey-specific feeding rates would also support this idea. Second, we ask what factors are driving within species-pair variation in feeding rates over space and time? To address this question we quantify variation in space and time for each interaction individually and determine the extent to which changes in prey density, conspecific predator density, prey body mass, or abiotic factors can explain this variation. Our results show that, despite considerable within-pair variation in feeding rates, ‘strong’ interactions tend to remain ‘strong’ while ‘weak’ interactions tend to remain ‘weak’. As a result, interaction strengths distributions show consistency in space and time. Our results also show that while prey density is a primary factor driving within-pair variation in feeding rates for many prey taxa, factors including prey body mass, water temperature, and stream discharge frequently exhibit even greater effects for other taxa.

## Methods

### Study sites

We estimated feeding rates of reticulate sculpin (*Cottus perplexus*) on its macroinvertebrate prey at nine stream sites within Oregon State University’s McDonald-Dunn Research Forest northwest of Corvallis, Oregon. The nine sites were each ~45 m in length and were nested within three streams (Berry, Oak, and Soap Creeks, see Preston et al. 2018). The three study streams were > 4km apart from one another, and the sites within each stream were on average 336 m apart (min = 87 m, max = 950 m). The ecology of streams in the McDonald-Dunn Research Forest has been well studied, including extensive work on the diverse (>325 species) macroinvertebrate community (e.g., Anderson and Lehmkuhl 1968, Kerst and Anderson 1975, Grafius and Anderson 1979), community interactions (e.g., Davis and Warren 1965, Hawkins and Furnish 1987), and ecosystem functioning (e.g. Warren et al. 1964). In addition to reticulate sculpin, other resident aquatic vertebrates at our sites include cutthroat trout (*Oncorhyncus clarkii*), Pacific giant salamanders (*Dicamptodon tenebrosus*), and brook lamprey (*Lampetra richardsoni*).

### Estimating feeding rates

We estimated feeding rates in an *in situ* manner by combining gut contents data from field surveys with information on the time period over which prey items remain identifiable in a sculpin’s stomach (hereafter the ‘prey identification time’). Prey-specific sculpin feeding rates were estimated for each survey as 
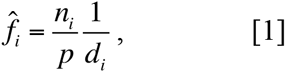

where *f̂_i_* is the average sculpin’s feeding rate (prey consumed predator^-1^ time^-1^), *n_i_* is the number of prey items of species *i* found in a sample of *p* predator stomachs, and *d_i_* is prey *i*’s estimated identification time (see also Novak and Wootton 2008, Novak et al. 2017, Wolf et al. 2017 and Preston et al. 2018). This approach explicitly accounts for differences in the amount of time that prey items persist in stomach contents, which when unaccounted for, will bias inferences about trophic interactions made from diet data (Hyslop 1980, Fairweather and Underwood 1983, Novak 2010, Preston et al. 2017).

### Field surveys

To collect sculpin diet information (*n_i_* and *p* in eqn. 1), we surveyed each of the nine stream sites in summer (June/July 2015), fall (September 2015), and spring (April 2016) (27 total site-by-season replicates). Sculpin were surveyed systematically throughout the whole area of the reach by a crew of four researchers using a backpack electroshocker (Smith-Root LR20B), a block net (1.0 x 1.0 m) and two dip nets (0.30 x 0.25 m). Block nets at each end prevented movement in and out of the reach during surveys. We anesthetized, weighed, measured, and nonlethally lavaged each sculpin with a 60 cc syringe and blunt 18-gauge needle to obtain gut contents. Sculpin were then held in aerated stream water and released after recovery from anesthesia. We preserved stomach contents in 70% ethanol and in the laboratory identified and measured prey for total body length. To estimate dry mass, we used a conversion factor based on wet mass for sculpin (Lantry and O’Gorman 2007) and length-to-mass regressions for invertebrates (Table S1). At each site, we also estimated prey densities by collecting macroinvertebrates with ten Surber samples (0.093 m^2^ in area) evenly spaced along the length of each reach. Surber samples were preserved in 70% ethanol and invertebrates were measured for body length and identified using Merritt et al. 2008. We quantified abiotic variables at each site, including stream discharge, canopy openness, substrate size, water temperature, and stream width (Supplemental Materials). Lastly, we estimated sculpin densities by correcting our electroshock sculpin counts using catch efficiency estimates from habitat-specific (pool or riffle) mark-recapture surveys conducted at each stream (Supplemental Materials).

### Prey identification times

Our estimates of prey identification times (*d_i_* in eqn. 1) were based on functions from laboratory trials during which individual sculpin were fed invertebrate prey and then lavaged over time to determine the rate at which prey became unidentifiable as a function of covariates. Our approach for estimating prey identification times is provided in detail in Preston et al. (2017) and is summarized in Preston et al. (2018). Here we provide an overview.

We estimated the prey-specific identification times for common prey types observed in sculpin diets, including mayflies (Ephemeroptera), stoneflies (Plecoptera), caddisflies (Trichoptera), flies (Diptera), beetles (Coleoptera), worms (Annelida), and snails (*Juga plicifera*) (Table S2). Our approach therefore incorporated differences in prey traits across taxonomic groups that affect rates of digestion by sculpin. In the laboratory trials, we varied water temperature (10°C to 20°C), prey size for each taxon (Table S2), and predator size (32 mm to 86 mm sculpin) in a continuous and randomized manner, and then fit Weibull survival curves to the observed prey status (identifiable or not) as a function of the covariates (Klein and Moeschberger 2005). The time periods over which sculpin were lavaged after feeding ranged from 10 min to 100 hrs depending on the prey type. The estimated laboratory coefficients from the Weibull survival functions were used with observed covariate information from our field surveys (i.e., predator and prey sizes and water temperatures) to estimate prey identification times for each prey item recovered from a sculpin’s stomach. For each prey item, the identification time was estimated as the mean of the probability density function that corresponded to the Weibull survival function under the observed covariate values (Preston et al. 2017). We then used the average prey-specific detection times within each survey to calculate the prey-specific sculpin feeding rates using eqn. 1. For prey types other than the aforementioned seven taxa, we used survival function coefficients from morphologically similar prey types (Table S3).

### Analyses

We first assessed changes in the overall distribution of all feeding rates in each survey by examining the distribution parameters including the mean, standard deviation, skewness and kurtosis. We then quantified the within-pair variation in feeding rates seen across space and time and compared it to the variation in feeding rates seen across species pairs at a given site and time. Within-pair variation was quantified as the coefficient of variation for each species pair using the mean and standard deviation of the prey-specific feeding rates across surveys. Not all prey taxa were observed in sculpin diets from all surveys, hence these calculations included up to 9 sites x 3 seasons = 27 feeding rate estimates for each species pair (Table S3). Across-pair variation was quantified for each survey as the coefficient of variation calculated using the mean and standard deviation of the survey’s prey-specific feeding rates. To quantify variation within and across species pairs we focused on the 25 prey taxa (i.e., pairwise interactions with sculpin) for which we had at least two feeding rate estimates per season and at least 10 estimates total across all site-season combinations (mean = 20.7 estimates; Table S3). Together, these taxa represented 88% of the individual prey items that we recovered (see *Results*).

Next we evaluated the consistency in the rank order of prey-specific feeding rates using Spearman’s correlation coefficients. We did this by ordering the 25 focal feeding rates by their overall means across all surveys and assessing deviations from this ordering in each of the individual surveys (n = 27 surveys). We also assessed deviations in the rank order across seasons (n = 3 seasons) and streams (n = 3 streams) using their respective mean values.

We examined the relative roles of space and time in contributing to the variation seen within each species pair (n=25) using a generalized linear mixed model (GLMM) with log-transformed feeding rates as the response (Zuur et al. 2009). Our model included the fixed effects of reach identity (i.e. ‘space’) and of season (i.e. ‘time’), and random intercept terms for stream (three reaches per stream) and prey taxonomic identity (up to 27 feeding rates per prey taxon). Diagnostic plots and comparisons to a model without random intercept terms indicated that inclusion of the random effects was justified (Supplemental Materials). To assess the contributions of ‘space’ and ‘time’ fixed effects, we compared the full model to 1) a model with season only, 2) a model with reach identity only, and 3) an intercept-only null model. We compared model performance using small sample size adjusted Akaike Information Criterion scores (AICc) (Burnham and Anderson 2002) and evaluated model fit using marginal and conditional R-squared values (Nakagawa and Schielzeth, 2013); marginal R-squared represents variance explained by fixed effects and conditional R-squared represents variance explained by fixed and random effects. To further examine feeding rate variation in space and time, we also calculated coefficients of variation for feeding rates on each of the 25 focal taxa across the nine sites (using mean feeding rates per site) and across the three seasons (using mean feeding rates per season).

Our next goal was to assess the capacity of prey density, predator density, prey body mass, and abiotic factors to explain the variation in prey-specific feeding rates we observed over space and time. These analyses focused on 20 of the 25 previously-considered prey taxa as 5 taxa (Dixidae, Ceratopogonidae, Copepoda, Ostracoda, Polycentropodidae) were sometimes not detected in surber samples, precluding estimates of their density. The analyses entailed using general linear models for each focal taxon, including a full model (all four hypothesized drivers included), models with each of the four predictors alone, and an intercept-only null model (six total models per prey taxon). While many other biologically reasonable models (i.e., variable combinations) are plausible, our primary goal was to determine the univariate explanatory power of the four variables rather than develop a predictive mechanistic model. Exploratory models also included a random intercept term for reach identity nested within stream, but these decreased relative model performance (based on AICc scores) and were thus not included in the final analysis (Supplemental Materials). Abiotic factors were incorporated as the first principal component from a PCA analysis of stream discharge, canopy openness, substrate size, water temperature, and stream width, using mean values per survey. Prey masses were from the Surber data and not the sculpin diet data. When a prey taxon was not detected in the Surber samples of a given survey, the corresponding feeding rate was omitted from the analysis. Feeding rates and all predictor variables other than the PC scores were log-transformed to improve conformance to model assumptions. For each prey taxon, we used AICc and R^2^ values to compare the six models. Lastly, we examined the overall univariate explanatory power of each of the four predictor variables across all taxa combined by summing the AICc scores of each variable’s prey-specific models. Plots showing covariate correlations and model residuals are shown in the Supplemental Materials (Figs. S1, S2).

## Results

### Feeding rate variation

The frequency distributions of feeding rates were positively skewed in all seasons and at all sites, exhibiting a consistent pattern of a few strong and many weak interactions (Fig. 1 and Fig. S3). Estimates of distribution skewness ranged from 1.4 to 5.4 (mean = 3.7) across surveys. These and the other distribution moments we measured did not differ consistently across streams or reaches, but did show seasonal differences in that all were generally highest in the summer (Table S4).

**Figure 1.**
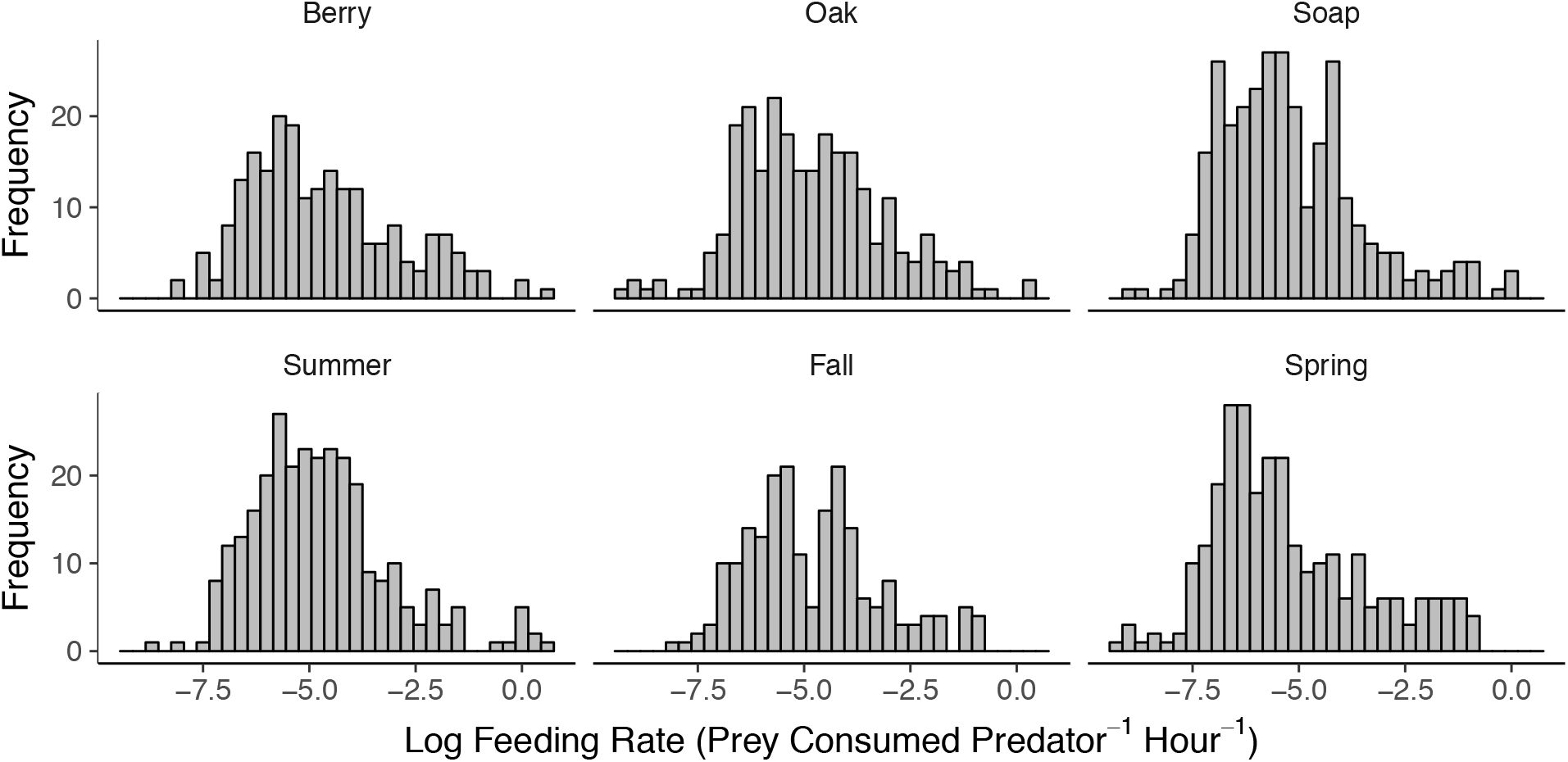
Log-transformed sculpin feeding rate distributions across three streams (top) and three seasons (bottom). Each distribution includes feeding rates from nine replicate surveys and all observed prey.

In total, we collected 15,471 identifiable prey items from 2,068 sampled sculpin. The 25 focal prey taxa accounted for 13,564 prey items (88% of the total). The majority of these focal prey items belonged to the orders Ephemeroptera (45%), Diptera (37%), Trichoptera (9%), and Plecoptera (5%). Mean prey-specific feeding rates across the focal taxa varied by over three orders of magnitude, with the highest mean feeding rates being on Baetidae mayflies and Chironomidae midges, and the lowest being on *Juga* snails (Fig. 2).

**Figure 2.**
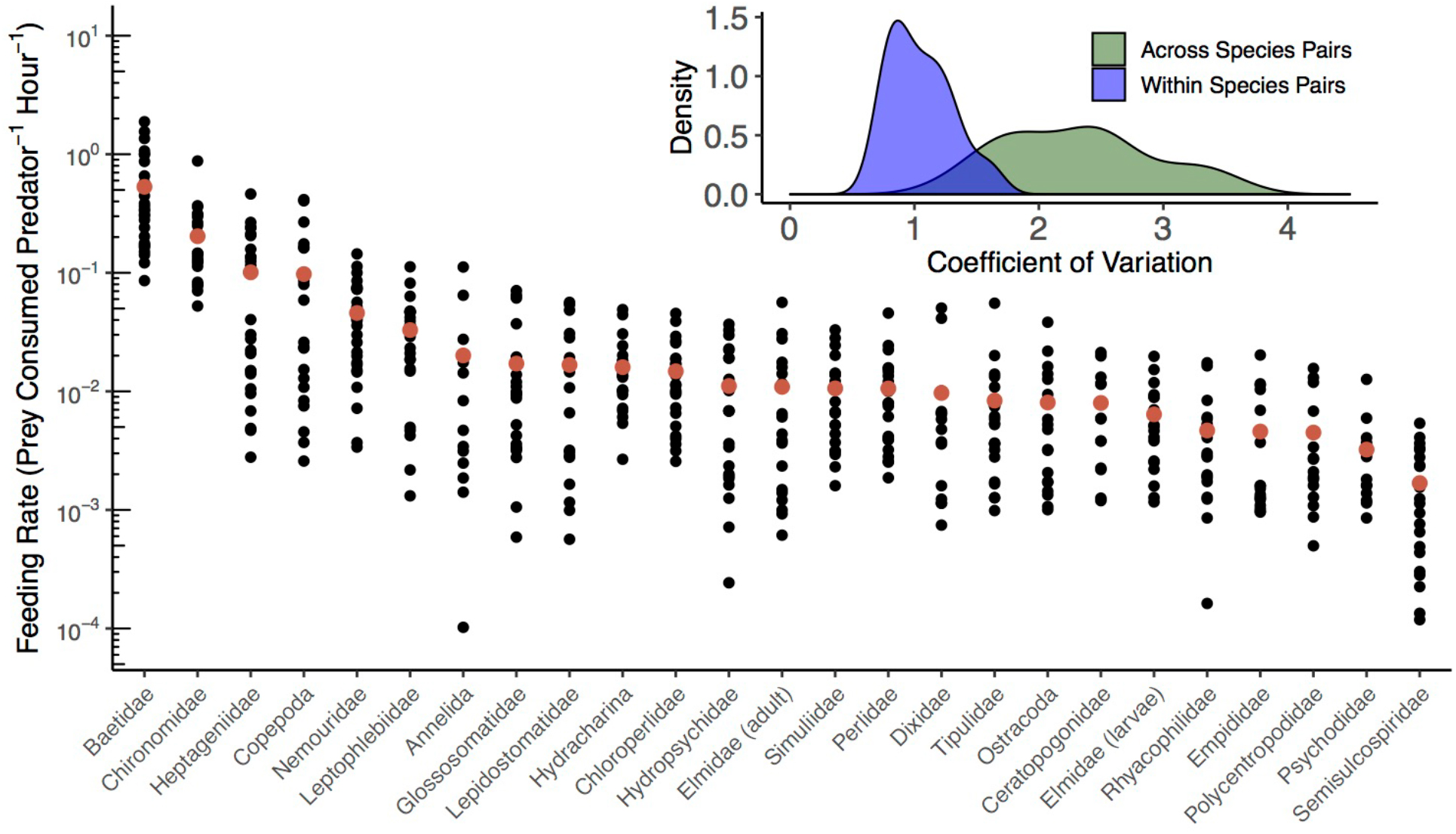
Feeding rates of reticulate sculpin on 25 taxa of invertebrate prey. In the main panel, the red points indicate the mean feeding rates across all surveys within a prey taxon and the black points represent individual surveys. Note the log-scale on the y-axis and that the prey-specific feeding rates are ordered by their means. The inset panel shows the replicate coefficients of variation for sculpin feeding rates within a species pair across surveys in space and time (n = 25 taxa shown in blue) and across species pairs within a survey at a specific site (n = 27 surveys shown in green). The mean CV within species pairs is 1.1 and the mean CV across species pairs is 2.3. In the larger panel, within pair variation corresponds to variation in the vertical direction for each prey taxon, while across pair variation corresponds to variation in the horizontal direction across prey taxa in a survey.

Overall, the variation in feeding rates across species pairs within a survey was greater than the variation across surveys within a species pair (Fig. 2 inset). The mean coefficient of variation was 2.31 across species pairs (min = 1.27, max = 3.53, median = 2.38; n=27 surveys), versus 1.05 for variation within species pairs (min = 0.71, max = 1.63, median = 1.01; n = 25 prey taxa). The within-pair difference from the lowest to highest feeding rates across all surveys in space and time averaged a 102-fold increase, ranging from 14-fold (Psychodidae flies) to 1093-fold (Annelid worms).

The rank order of prey-specific feeding rates remained relatively consistent across seasons, streams, and individual surveys (Fig. 2 and Fig. S4). The ordering of mean feeding rates across the three seasons (ρ = 0.92 in summer; 0.84 in fall; 0.80 in spring) and the three streams (ρ = 0.86 at Berry Creek; 0.91 at Oak Creek; 0.86 at Soap Creek) did not differ greatly from the ordering of the overall means (Fig. S4). Across surveys the mean Spearman’s correlation coefficient was 0.71 (range = 0.45 to 0.93), with deviations from the order of the mean feeding rates driven primarily by variation in the lowest feeding rates.

### Effects of space and time on within-pair variation

Many prey-specific feeding rates showed strong seasonal variation. Summer corresponded to the highest feeding rates for 17 of the 25 prey taxa, followed by spring (7 taxa), and fall (1 taxon) (Fig. 3). Feeding rates on flies (Figs 3a to 3g) and worms (Fig. 3t) showed relatively little seasonal change, while mayflies (Figs. 3h to 3j), stoneflies (Figs. 3k to 3m), and caddisflies (Figs. 3n to 3r) showed larger seasonal differences. Among the largest seasonal changes in mean feeding rates were those observed on mayflies, including Baetidae (a 4-fold decrease in mean feeding rates from summer to fall; Fig. 3h) and Heptageniidae (a 10-fold increase in mean feeding rates from fall to spring; Fig. 3i). General linear models fit to all 25 prey-specific feeding rates combined supported the idea that feeding rate variation was more strongly associated with season than reach identity (Table S5). The top-performing model included season alone. Nevertheless, even in the top-performing model, the fixed effect of season explained relatively little variation in feeding rates (marginal R^2^ = 0.04) compared to the random effect of prey taxon (conditional R^2^ = 0.63).

**Figure 3.**
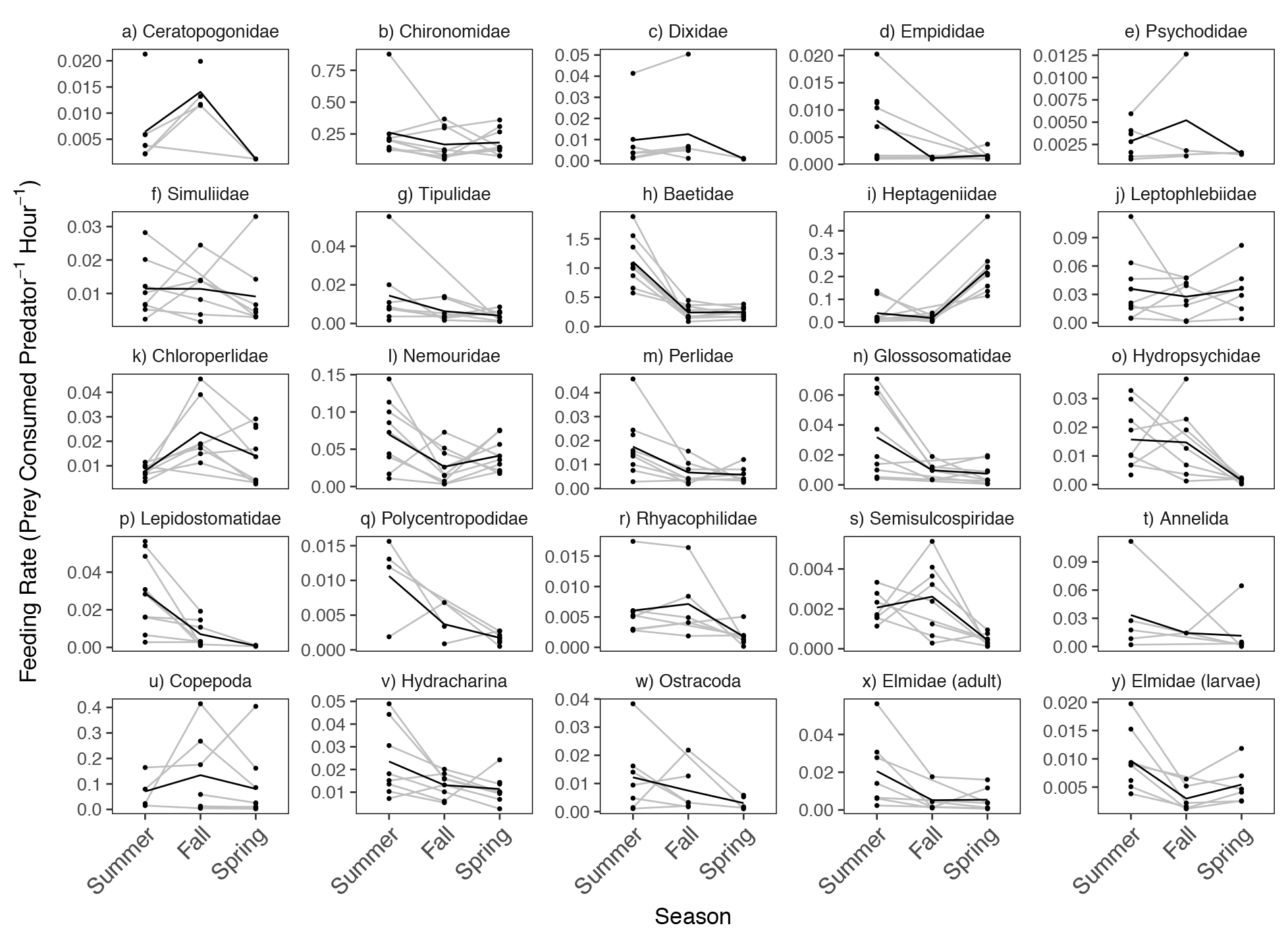
Spatial and temporal variation in reticulate sculpin feeding rates on 25 invertebrate prey taxa. The black lines show the seasonal mean feeding rates and the grey lines connect the same sites over time. Note the differences in the y-axis scale across panels.

The coefficients of variation for feeding rates in space versus time reflected the different effects of season on each prey taxon. The CVs were higher across sites than across seasons for 7 of 8 fly and worm taxa (Fig. S5). In contrast, for mayflies, stoneflies, and caddisflies, the CVs were higher across seasons than across sites for 8 of 11 taxa (Fig. S5). In general, the taxa with high variation across seasons showed consistent differences in mean seasonal feeding rates (Fig. 3), whereas taxa showing higher variation in space were not necessarily associated with consistent differences in mean reach- or stream-level feeding rates.

### Drivers of within-pair variation

The four hypothesized explanatory variables for within-pair variation in feeding rates (i.e., prey density, prey body mass, predator density, and abiotic factors) varied more across seasons than across sites for most prey taxa. The densities for 9 of the 20 taxa considered in these prey-specific analyses were highest in summer, while another 9 were highest in fall and two were highest in spring (Fig. S6). Nine of the 20 taxa had the largest mean body size in spring (Fig. S7). Of the abiotic variables measured, water temperature and stream discharge showed the largest variation, with low flows (mean = 0.01 m^3^s^1^) and warm temperature (mean =15°C) in summer, followed by lower temperatures (mean =10°C) and higher flows (mean = 0.09 m^3^s^-1^) in spring (Fig. S8). The first principal component from the PCA analysis, which was associated with 41% of the variation in the abiotic data, was positively associated with lower water temperatures and higher discharge (Fig. S9). Sculpin densities were highest in summer (mean = 2.8 m^-2^) and decreased slightly in fall and spring (mean = 2.1 m^-2^ for both) (Fig. S10).

Variation in prey density and abiotic factors showed relatively consistent directional associations with within-pair variation in feeding rates. Feeding rates increased with prey density for 18 of the 20 prey taxa (Fig. 4, Table S6); the two exceptions being Empididae flies and Hydracharina mites which showed negative relationships. The first principal component of our PCA analysis of abiotic variables was negatively associated with feeding rates for 14 of the 20 taxa (Fig. 6, Table S6), indicating that feeding rates decreased at lower temperatures and higher flows.

**Figure 4.**
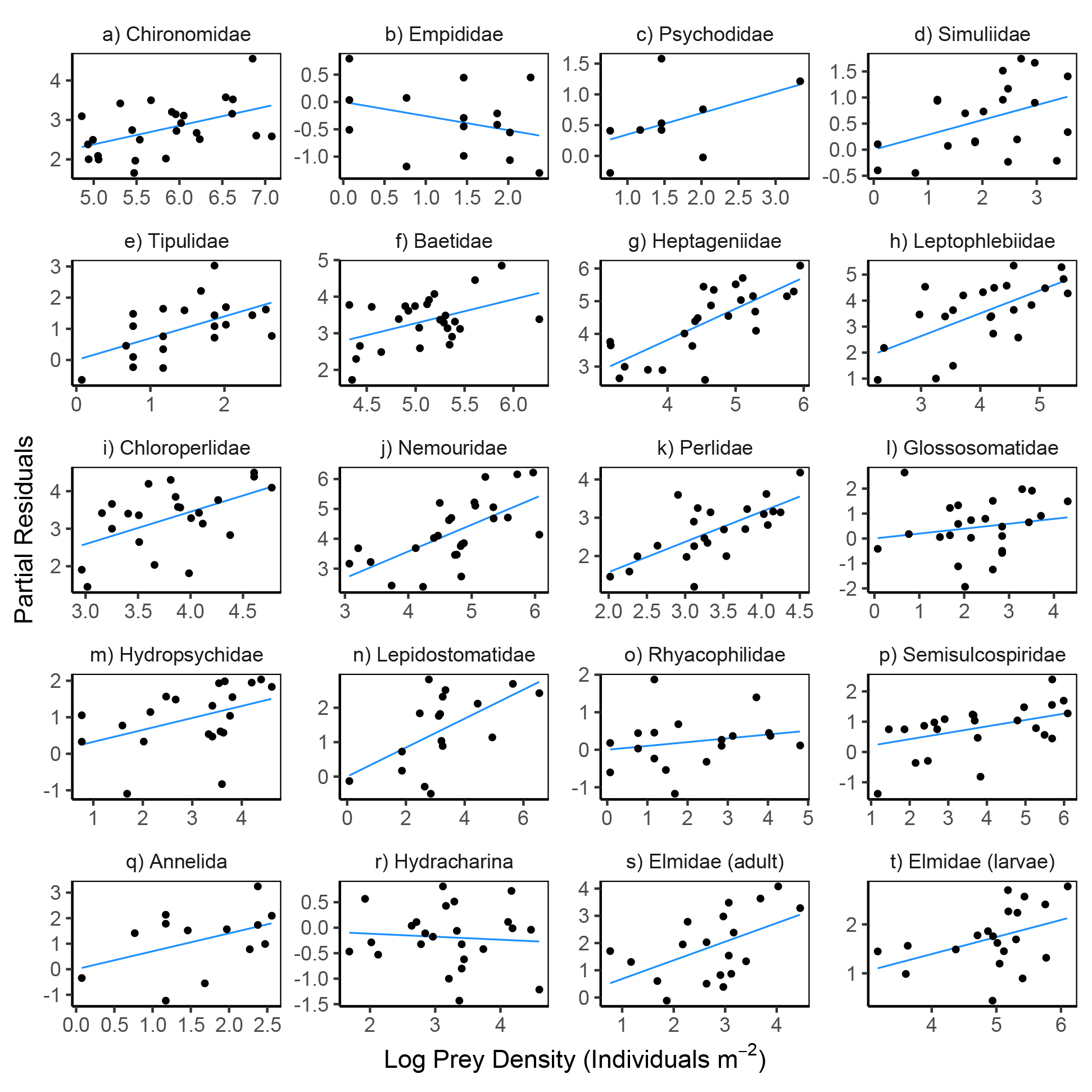
Partial residual plots for models predicting reticulate sculpin feeding rates on 20 prey taxa. The regression lines show partial fits from the prey-specific full models (Table S6) which included prey density, prey body mass, predator density, and abiotic factors. The points show the full model residuals + *β_i_X_i_*, where *β_i_* is the regression coefficient for prey density from the full model and *X_i_* is the observed prey density shown on the x-axis. Note the varying scales for log prey density across panels.

**Figure 5.**
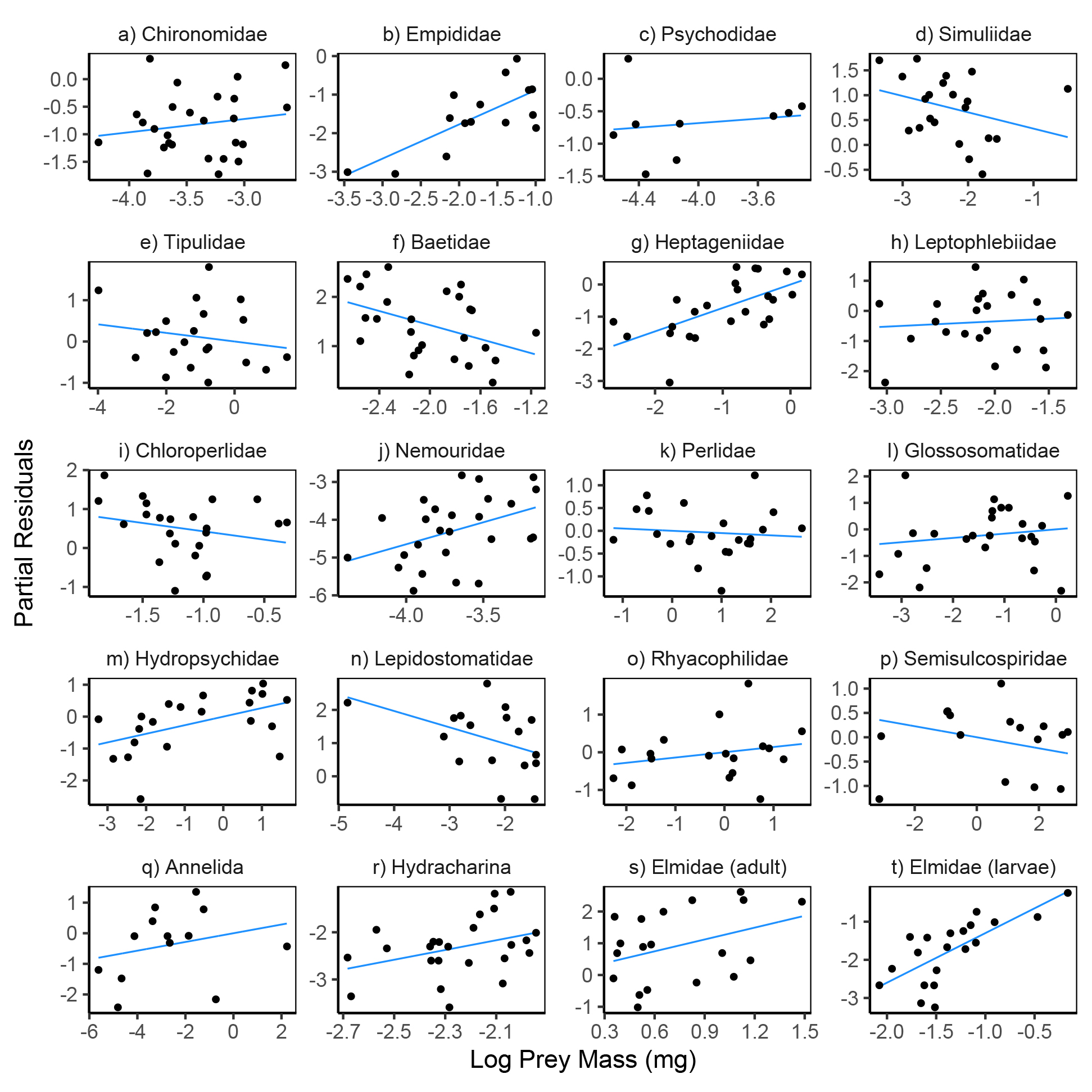
Partial residual plots for models predicting reticulate sculpin feeding rates on 20 prey taxa. The regression lines show partial fits from the prey-specific full models (Table S6) which included prey density, prey body mass, predator density, and abiotic factors. The points show the full model residuals + *β_i_X_i_*, where *β_i_* is the regression coefficient for prey mass from the full model and *X_i_* is the observed prey mass shown on the x-axis. Note the varying scales for log prey mass across panels.

**Figure 6.**
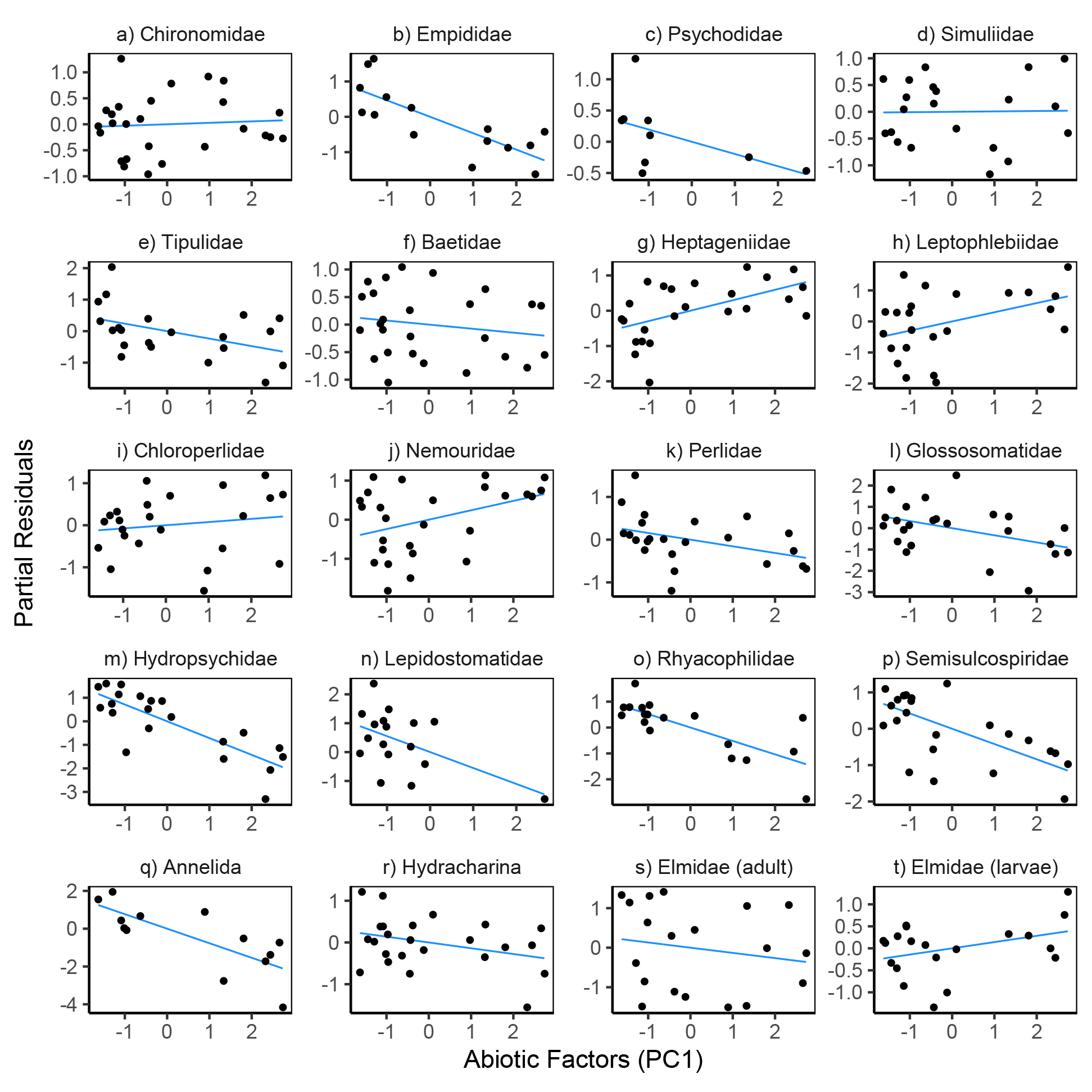
Partial residual plots for models predicting reticulate sculpin feeding rates on 20 prey taxa. The regression lines show partial fits from the prey-specific full models (Table S6) which included prey density, prey body mass, predator density, and abiotic factors. The points show the full model residuals + *β_i_X_i_*, where *β_i_* is the regression coefficient for the abiotic factors from the full model and *X_i_* is the first principal component from a PCA analysis of the abiotic factors.

The directional nature of the relationships between feeding rates and variation in prey mass and sculpin density differed widely across the 20 taxa. Prey body mass was positively associated with feeding rates for 13 taxa and negatively associated for 7 taxa, without a clear taxonomic divide in either the sign or magnitudes of correlations (Fig. 5, Table S6). Sculpin densities were positively associated with feeding rates for half of the taxa and negatively associated with feeding rates for the other half (Fig. 7, Table S6).

**Figure 7.**
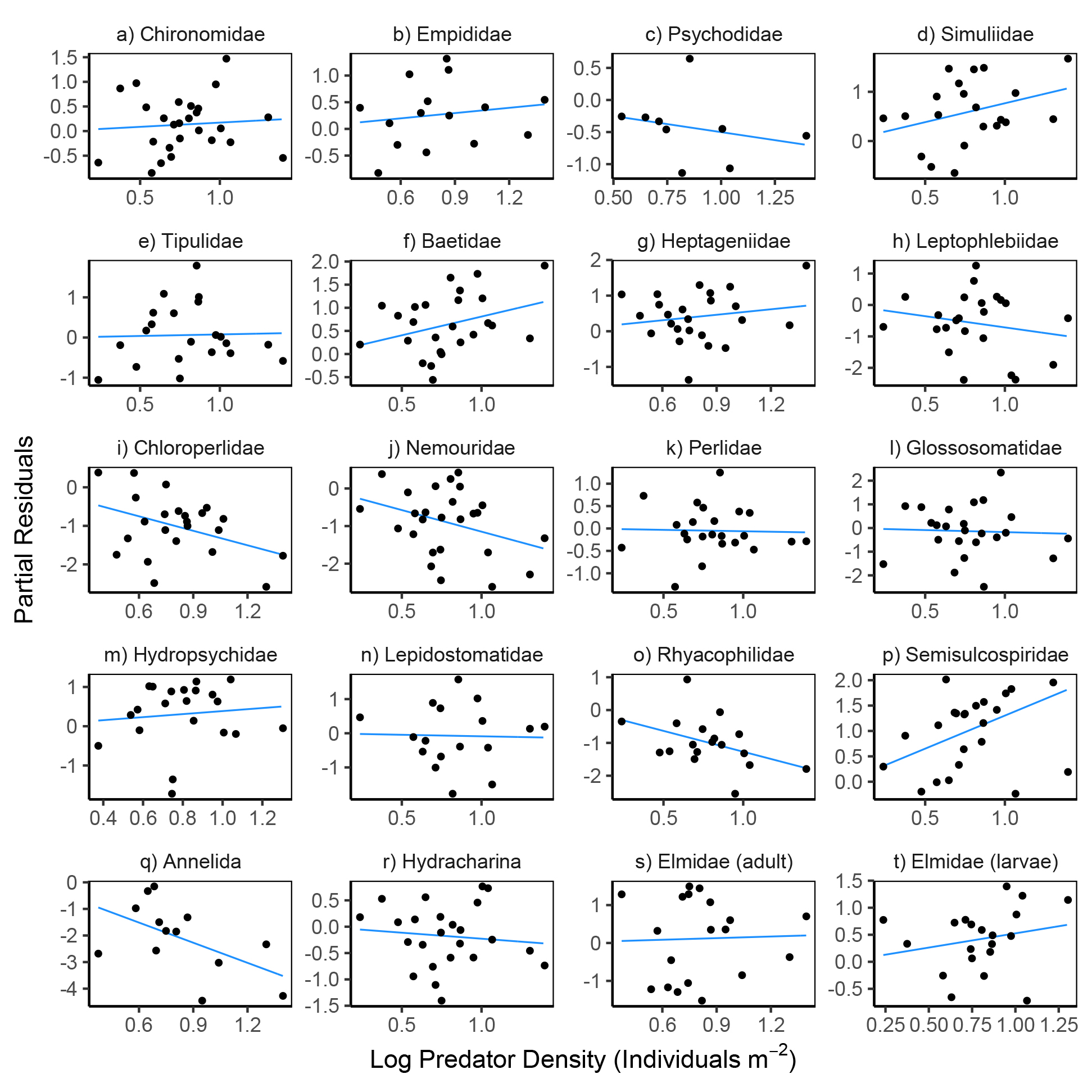
Partial residual plots for models predicting reticulate sculpin feeding rates on 20 prey taxa. The regression lines show partial fits from the prey-specific full models (Table S6) which included prey density, prey body mass, predator density, and abiotic factors. The points show the full model residuals + *β_i_X_i_*, where *β_i_* is the regression coefficient for predator (sculpin) density from the full model and *X_i_* is the observed predator density shown on the x-axis.

The ability of prey density, prey body mass, predator density, and abiotic factors to explain variation in feeding rates differed widely across the 20 prey taxa (Table S7). For 3 taxa (Baetidae, Hetpageniidae, Semisculcospiridae), the top-performing model was the full model with all four covariates. For 2 taxa (Psychodidae and Chloroperlidae), the intercept-only model outperformed all other models. Of the other 15 taxa, the top model included only prey density for 7 taxa (Elmidae adults, Chironomidae, Tipulidae, Leptophlebidae, Nemouridae, Perlidae, Lepidostomatidae), abiotic factors for 5 taxa (Empididae, Annelida, Glossosomatidae, Hydropsychidae, Rhyacophilidae), and prey mass for 3 taxa (Hydracharina, Elmidae larvae, Simulidae). The variation explained by the top models averaged 37% (R^2^ range: 0.14 to 0.81) after excluding the two taxa for which the intercept-only model performed best (Table S7). Summing the AICc scores across the 20 taxa resulted in the model including only abiotic factors having the best relative performance, followed by models including only prey density (∆AICc = 6.1), prey mass (∆AICc = 47.1), the intercept-only model (∆AICc = 66.7), the full model (∆AICc = 90.2), and only predator density (∆AICc = 98.9).

## Discussion

Ecologists are increasingly grappling with the dynamic nature of species interactions and the consequences of food web variation for the structure and functioning of communities (Poisot et al. 2015, Lopez et al. 2017, Tylianakis and Morris 2017). Alongside this growing appreciation for variation, there remains the now longstanding realization that many properties of communities are conserved. This includes the recurrently observed skewed distribution of interaction strengths (many weak, few strong), which has often been linked to the stability of species-rich communities (McCann et al. 1998, Borrvall et al. 2000, Wootton and Emmerson 2005, Gellner and McCann 2016). This leads to an interesting inconsistency: if interactions in food webs are highly dynamic, why is the skewed distribution of interaction strengths apparently conserved? Our results help to reconcile the seemingly contradictory nature of these two food web properties by showing how the ecological scale of inference shapes conclusions about whether species interactions are dynamic or consistent. We show that, from the perspective of community-wide variation in interaction strength, the pairwise interactions in our study system are relatively consistent in space and time; ‘strong’ interactions remain ‘strong’ and ‘weak’ interactions remain ‘weak’. Although within-pair variation averaged a 100-fold difference from the lowest to highest observed feeding rate, it was much smaller than the variation seen across pairs, leading to a consistent rank order and overall frequency distribution of species interaction strengths. Our results therefore emphasize how food webs encompassing dynamic interactions can yield properties that are nonetheless conserved in space and time.

The consistency of the pairwise interactions relative to community-wide variation has several implications. Among these is that it suggests the existence of fundamental characteristics that drive interaction strengths within versus between interacting species pairs. That is, while within-pair variation was explained by distinct prey-specific factors (especially abiotic variables and prey density), the variation seen across species pairs suggests a more fundamental role of prey identity in driving variation (Preston et al. 2018). Clearly, species identity is associated with a wide range of attributes of relevance to foraging predators (e.g., life history traits) that will have shown consistent differences across prey taxa over the spatiotemporal scale of our surveys. The relative consistency of the pairwise interactions also provides mechanistic insight into processes underlying other conserved food web properties beyond the skewed distribution of interaction strengths. For instance, the persistence of common structural characteristics in food webs, such as motif frequencies and “backbone” interactions (Mora et al. 2018), may be explained by the relative consistency of species interaction strengths at the community scale. Thus, while our findings should be interpreted within the spatial, temporal, and ecological (i.e., focal prey community) scale of our surveys, they collectively suggest that from a community-wide perspective, interaction strengths may be more predictable than commonly assumed.

The relative consistency of the pairwise interactions seen in our study is nonetheless striking given that we focused on a generalist predator in species-rich streams that typically show large variation in biotic and abiotic factors (Fisher et al. 1982, Power et al. 2008, Death 2010). Pairwise interaction strengths in streams are expected to be dynamic because interaction-strength altering abiotic drivers themselves vary greatly over time (Power et al. 1988, Peckarsky et al. 1990, Wootton et al. 1996, Tonkin et al. 2017). Spatial heterogeneity in stream habitat can also shape community structure over small spatial scales (Palmer and Poff 1997), and the life-histories of many stream organisms result in fluctuating population abundances and size structures across seasons (Huryn and Wallace 2000). The observed variation in prey-specific feeding rates in our study is usefully interpreted in the context of these factors.

When considered in a univariate fashion, abiotic factors explained the most variation in feeding rates for five prey taxa, suggesting that knowledge traditionally seen as vital to the nature of predator functional responses was of little utility for these species. The importance of abiotic conditions was further underscored by the result that the prey-specific models including only abiotic factors had the lowest total score when their AICc scores were summed across the 20 taxa that this analysis included. This role of abiotic factors in driving feeding rate variation was less apparent in our previous single-season study (Preston et al. 2018), which emphasizes the importance of spatiotemporal replication and scale-dependence in considering interaction strength variation and its drivers. Here, we observed a 33% decrease in water temperature and a 9-fold increase in stream flows from summer to the following spring, which far exceeded the abiotic variation in our previous study. Prey-specific sculpin feeding rates correlated negatively with stream flow and positively with water temperatures, consistent with expected effects of these variables on energetic demands, activity levels, and possibly foraging conditions for fishes (e.g., ability to locate and consume prey under high flow and low water clarity) (Elwood and Waters 1969, Kishi et al. 2005). These relationships with abiotic variables were particularly strong for feeding rates on caddisflies. More broadly, the observed role of abiotic factors in our study supports the idea that predation should be stronger (and more consistent) in low stress environments and weaker in high stress environments (Peckarsky 1983, Menge and Sutherland 1987, Peckarsky 1990).

We found that univariate prey-density models had the second best performance behind models with abiotic variables in explaining within-pair feeding rate variation. For most prey (18 of 20 taxa), prey-specific feeding rates increased as prey-specific density increased, suggesting that sculpin are opportunistically consuming prey that they encounter, especially at low prey densities. The relative role of prey density across pairwise interactions was associated with the life histories of the prey taxa, such as voltinism and length of the nymphal period. These factors also contributed to differences in variation associated with space versus time. In general, the seasonal patterns in feeding rates on mayflies, stoneflies, and caddisflies were likely driven by their mostly univoltine lifecycles, where densities and size distributions change markedly over the season and often peak in spring and summer (Anderson and Wold 1972, Kerst and Anderson 1974). For instance, the large seasonal changes in feeding rates on Baetid mayflies (highest in summer) and Heptageniid mayflies (highest in spring) corresponds with peak emergence periods, after which decreases in nymphal densities due to emergence result in lower feeding rates (Lehmkuhl 1968, 1969). Feeding rates on prey taxa that showed less seasonal variation may in turn be related to longer larval periods or multiple generations per year that result in less temporal fluctuation in prey density and size (e.g., many dipterans; Dudley and Anderson 1987).

The slopes of the within-pair relationships between feeding rates and the densities of each prey taxon are informative because they allow comparisons with predictions from predator functional response models. These slopes were less than one on the log-log scale for all but one prey taxon (for which the slope was approximately one)(Table S6) reflecting decelerating positive (i.e., saturating) relationships on the natural scale (Menge et al. 2018). This finding is consistent with nearly all parametric models of predator functional responses (Jeschke et al. 2002). The mechanisms underlying the saturation of prey-specific feeding rates, however, are not necessarily clear in that an accelerating (non-saturating) slope between feeding rates and prey densities was observed when considering the relationship across all prey species combined (Preston et al. 2018 and Fig. S2). In other words, feeding rates increased with within-species differences in prey density in a decelerating (saturating) form but increased with between-species differences in prey density in an accelerating (non-saturating) fashion. Further lines of evidence also suggest that the overall feeding rates of sculpin are not limited by either handling or digestion times, which are the typically invoked rate-limited steps for generating saturating functional responses. For instance, the mean number of prey observed per sculpin was less than 30-times the maximum observed, suggesting that most sculpin are able to consume far more prey than is observed in their stomachs (a widespread characteristic of fishes; Armstrong and Schindler 2011). It is also noteworthy that for most taxa (13 of 20) the top-performing model did not include prey density, and that for 8 taxa prey body mass or abiotic factors explained more of the univariate variation in within-pair feeding rates. Together, these findings suggest that functional response models that focus only on within-prey variation in density may poorly predict feeding rates in the field, and that current functional response models (developed on the basis of within-prey variation) may not be as easily scaled-up to predicting total or between-prey variation in feeding rates as currently assumed. Future work is needed to develop and test functional response models for species-rich contexts that can better characterize and assess the interdependencies between the prey-specific feeding rates of generalist predators (Abrams 2001, Novak et al. 2017).

Prey mass was most closely associated with feeding rate variation for relatively few prey taxa (3 of 20), suggesting that efforts to infer interaction strengths based on pairwise predator-prey size relationships should be applied to food webs containing generalist predators with caution. Across the entire feeding rates dataset, there appears to be an ‘optimal’ prey mass associated with the highest feeding rates (Preston et al. 2018 and Fig. S2). In general, predators are thought to select for prey of intermediate predator-prey body size ratios, thereby increasing energetic gains from prey while avoiding large prey that are less efficiently consumed (Brose 2010, Kalinkat et al. 2013). This could result in either monotonic positive or negative relationships between feeding rates and prey size within a given prey taxon depending on where a prey type lies relative the optimum. We observed both positive and negative correlations between prey mass and prey-specific feeding rates in our analysis, but the direction of the relationships were not consistent with a single optimal prey mass across all prey taxa. Some prey likely showed negative relationships between mass and feeding rates because large prey individuals present challenges for consuming and digesting prey. For instance, limitations on sculpin gape width and digestibility likely play a role in the low feeding rates on large *Juga* snails (Semisulcospiridae) (Preston et al. 2017). For other feeding rates that showed strong relationships with prey size (e.g., Elmidae beetle larvae), feeding rates were highest when prey were largest. While for some taxa the specific mechanisms underlying this pattern are not clear, it is possible that prey size is correlated with other traits (e.g., anti-predator behaviors, propensity to drift) that could contribute to the different direction of correlations between size and feeding rates across taxa. Recent research also indicates that differences in the mean and the standard deviation of predator-prey size relationships across food webs are likely linked to changes in overall prey availability (Costa-Pereira et al. 2018). While we did not examine interactions between explanatory variable (e.g., prey density and prey mass), this provides an interesting area for future work.

Predator (i.e., sculpin) density was not a primary factor underlying variation in prey-specific feeding rates in our dataset. The presence and relative importance of predator dependence has been a debated topic in the literature (Abrams and Ginzburg 2000, Baraquand 2014), with relatively few studies having assessed predator dependence in field settings (Novak et al. 2017). The lack of a relationship for most prey taxa is interesting given that 1) we observed a negative correlation between sculpin density and feeding rates in the dataset across all combined prey taxa from summer (Preston et al. 2018), and 2) sculpin in streams are known to be territorial such that increases in density are expected to increase intraspecific interactions and decrease time spent feeding (Grossman et al. 2006). It is possible that wider variation in predator densities, beyond what was observed naturally at our sites would be more effective at revealing whether or not predator interference occurs in this system. That said, our results suggest that over the observed range of species densities predator dependence is unlikely to strongly shape sculpin feeding rates relative to other factors.

Several aspects of our study are of relevance in evaluating the generality of our results and the degree to which they can be extrapolated to other food webs. We focused on a single predator species and, for most analyses, only a subset of its prey taxa. While sculpins are generalists, as are most predators, focusing on other predators in our system could have altered some conclusions. For instance, our own preliminary evidence suggests that cutthroat trout in our streams show less consistency in their prey-specific feeding rates across seasons due to highly variable terrestrial prey availability (Falke et al. in prep). Similarly, although our focus on a subset of the sculpin prey taxa was necessitated by the lack of replicated feeding rate estimates for many prey taxa, consideration of the entire prey community would likely increase both within and across-pair variation in feeding rates. Furthermore, while streams in general are highly dynamic, our sites typically do not dry completely in late summer unlike some Mediterranean climate streams and they no longer support anadromous fishes, both of which can drive wholesale food web alternations (Gasith and Resh 1999, Naiman et al. 2002). Nonetheless, we do not expect any of these study system choices to have strongly affected our main conclusions.

Overall our results indicate that, while the complexity and dynamics of food webs can appear intractable (Polis 1991), at least some interaction attributes are more consistent than often recognized. As a result, the dynamics of trophic interactions may be predictable over space and time based on characteristics of the interacting species and their environment. Species interactions can thus be highly dynamic while still generating empirical patterns that prove ubiquitous across unique food webs (Wootton and Emmerson 2005). Promising next steps in efforts to understand and predict species interactions will require developing and testing mechanistic models that incorporate species densities, species traits beyond body size, and environmental covariates in shaping the strength and functional form of species interactions in species-rich communities. Achieving this aim will benefit from future empirical work that bridges across scales of interactions in space and time, ranging from species pairs to whole food webs.

## Acknowledgements (Note that Landon says there are other students that need to be added here)

For assistance with data collection we thank Madeleine Barrett, Alicen Billings, Daniel Gradison, Kurt Ingeman, Tamara Layden, Dana Moore, Arren Padgett, Wendy Saepharn, Alex Scharfstein, Johnny Schwartz, Leah Segui, Isaac Shepard, Samantha Sturman, Ernesto Vaca Jr, and Beatriz Werber. Richard Van Driesche and Michael Bogan provided input on identifying aquatic invertebrates and Stan Gregory provided background information on our field sites. We thank the Oregon State University College of Forestry for providing access to field sites in the McDonald-Dunn Research Forest. Funding was provided by the National Science Foundation Grant DEB-1353827, Oregon State University, and the University of Wisconsin-Madison. Data are available from the Dryad Digital Repository: http://(upon publication).

## References

Abrams, P. A. 2001. Describing and quantifying interspecific interactions: a commentary on recent approaches. Oikos 94:209–218.

Abrams, P. A., and L. R. Ginzburg. 2000. The nature of predation: prey dependent, ratio dependent or neither? Trends in Ecology and Evolution 15: 337-341.

Anderson, N. H., and D. M. Lehmkuhl. 1968. Catastrophic drift of insects in a woodland stream. Ecology 49: 198-206.

Anderson, N. H., and J. L. Wold. 1972. Emergence trap collections of Trichoptera from an Oregon stream. The Canadian Entomologist 104:189–201.

Armstrong, J. B., and D. E. Schindler. 2011. Excess digestive capacity in predators reflects a life of feast and famine. Nature 476:84-88.

Baraquand, F. 2014. Functional responses and predator-prey models: a critique of ratio dependence. Theoretical Ecology 7:3-20.

Bellmore, J. R., C. V. Baxter, and P. J. Connolly. 2015. Spatial complexity reduces interaction strengths in the meta-food web of a river floodplain mosaic. Ecology 96:274–283.

Borrvall, C., B. Ebenman, and T. J. Tomas Jonsson. 2000. Biodiversity lessens the risk of cascading extinction in model food webs. Ecology Letters 3:131–136.

Boutin, S., C. J. Krebs, R. Boonstra, M. R.T. Dale, S. J. Hannon, K. Martin, A. R.E. Sinclair, J. N.M. Smith, R. Turkington, and M. Blower. 1995. Population changes of the vertebrate community during a snowshoe hare cycle in Canada’s boreal forest. Oikos 74:69–80.

Brose, U. 2010. Body-mass constraints on foraging behaviour determine population and food-web dynamics. Functional Ecology 24:28–34.

Burnham, K. P., and D. R. Anderson. 2002. Model Selection and Multimodel Inference: a Practical Information-Theoretic Approach. Springer. New York, NY.

Byers, J. E., Z. C. Holmes, and J. C. Malek. 2017. Contrasting complexity of adjacent habitats influences the strength of cascading predatory effects. Oecologia 185:107–117.

Costa-Pereira, R., M. S. Araújo, R. da S. Olivier, F. L. Souza, and V. H. Rudolf. 2018. Prey limitation drives variation in allometric scaling of predator-prey interactions. The American Naturalist 192:139–149.

Cross, W. F., C. V. Baxter, E. J. Rosi-Marshall, R. O. Hall, T. A. Kennedy, K. C. Donner, W. Kelly, A. Holly, S. E. Seegert, K. E. Behn, and others. 2013. Food-web dynamics in a large river discontinuum. Ecological Monographs 83:311–337.

Davis, G. E., and C. E. Warren. 1965. Trophic relations of a sculpin in laboratory stream communities. The Journal of Wildlife Management 29:846–871.

Death, R. G. 2010. Disturbance and riverine benthic communities: what has it contributed to general ecological theory? River Research and Applications 26:15–25.

de Roos, A. M., L. Persson, and E. McCauley. 2003. The influence of size-dependent life-history traits on the structure and dynamics of populations and communities. Ecology Letters 6:473–487.

Dudley, T. L., and N. H. Anderson. 1987. The biology and life cycles of Lipsothrix spp. (Diptera: Tipulidae) inhabiting wood in Western Oregon streams. Freshwater Biology 17:437–451.

Elwood, J. W., and T. F. Waters. 1969. Effects of floods on food consumption and production rates of a stream brook trout population. Transactions of the American Fisheries Society 98:253–262.

Fagan, W. F., and L. E. Hurd. 1994. Hatch density variation of a generalist arthropod predator: population consequenes and community impact. Ecology 75:2022–2032.

Fairweather, P. G., and A. J. Underwood. 1983. The apparent diet of predators and biases due to different handling times of their prey. Oecologia 56:169–179.

Fisher, S. G., L. J. Gray, N. B. Grimm, and D. E. Busch. 1982. Temporal succession in a desert stream ecosystem following flash flooding. Ecological monographs 52:93–110.

Gasith, A., and V. H. Resh. 1999. Streams in Mediterranean climate regions: abiotic influences and biotic responses to predictable seasonal events. Annual Review of Ecology and Systematics 30:51–81.

Gellner, G., and K. S. McCann. 2016. Consistent role of weak and strong interactions in high- and low-diversity trophic food webs. Nature communications 7:11180.

Grafius, E., and N. H. Anderson. 1979. Population dynamics, bioenergetics, and role of *Lepidostoma quercina* Ross (Trichoptera: Lepidostomatidae) in an Oregon woodland stream. Ecology 60:433–441.

Grossman, G. D., R. E. Ratajczak, J. T. Petty, M. D. Hunter, J. T. Peterson, and G. Grenouillet. 2006. Population dynamics of mottled sculpin (Pisces) in a variable environment: information theoretic approaches. Ecological Monographs 76:217–234.

Hawkins, C. P., and J. K. Furnish. 1987. Are snails important competitors in stream ecosystems? Oikos 49:209–220.

Holling, C. S. 1959. Some characteristics of simple types of predation and parasitism. The Canadian Entomologist 91:385–398.

Huryn, A. D., and J. B. Wallace. 2000. Life history and production of stream insects. Annual Review of Entomology 45:83–110.

Hyslop, E. J. 1980. Stomach contents analysis—a review of methods and their application. Journal of Fish Biology 17:411–429.

Jeschke, J. M., M. Kopp, and R. Tollrian. 2002. Predator functional responses: discriminating between handling and digesting prey. Ecological Monographs 72:95–112.

Kalinkat, G., F. D. Schneider, C. Digel, C. Guill, B. C. Rall, and U. Brose. 2013. Body masses, functional responses and predator–prey stability. Ecology Letters 16:1126–1134.

Kalinoski, R. M., and J. P. DeLong. 2016. Beyond body mass: how prey traits improve predictions of functional response parameters. Oecologia 180:543–550.

Kerst, C. D., and N. H. Anderson. 1974. Emergence patterns of Plecoptera in a stream in Oregon, USA. Freshwater biology 4:205–212.

Kerst, C. D., and N. H. Anderson. 1975. The Plecoptera community of a small stream in Oregon, USA. Freshwater Biology 5:189–203.

Kishi, D., M. Murakami, S. Nakano, and K. Maekawa. 2005. Water temperature determines strength of top-down control in a stream food web. Freshwater Biology 50:1315–1322.

Klein, J. P., and M. L. Moeschberger. 2005. Survival analysis: techniques for censored and truncated data. Springer Science & Business Media. New York, NY.

Lantry, B. F., and R. O’Gorman. 2007. Drying temperature effects on fish dry mass measurements. Journal of Great Lakes Research 33:606–616.

Lehmkuhl, D. M. 1968. Observations on the life histories of four species of *Epeorus* in western Oregon. Pan-Pacif. Entomologist 44:129–137.

Lehmkuhl, D. M. 1969. Biology and downstream drift of some Oregon Ephemeroptera. PhD Dissertation. Oregon State University.

Lopez, D. N., P. A. Camus, N. Valdivia, and S. A. Estay. 2017. High temporal variability in the occurrence of consumer–resource interactions in ecological networks. Oikos 126:1699–1707.

McCann, K., A. Hastings, and G. R. Huxel. 1998. Weak trophic interactions and the balance of nature. Nature 395:794–798.

Menge, B. A. 2000. Top-down and bottom-up community regulation in marine rocky intertidal habitats. Journal of Experimental Marine Biology and Ecology 250:257–289.

Menge, B. A., E. L. Berlow, C. A. Blanchette, S. A. Navarrete, and S. B. Yamada. 1994. The keystone species concept: variation in interaction strength in a rocky intertidal habitat. Ecological Monographs 64:249–286.

Menge, B. A., and J. P. Sutherland. 1987. Community regulation: variation in disturbance, competition, and predation in relation to environmental stress and recruitment. The American Naturalist 130:730–757.

Menge, D. N., A. C. MacPherson, T. A. Bytnerowicz, A. W. Quebbeman, N. B. Schwartz, B. N. Taylor, and A. A. Wolf. 2018. Logarithmic scales in ecological data presentation may cause misinterpretation. Nature Ecology & Evolution 2:1393-1402.

Mora, B. B., D. Gravel, L. J. Gilarranz, T. Poisot, and D. B. Stouffer. 2018. Identifying a common backbone of interactions underlying food webs from different ecosystems. Nature Communications 9:2603.

Naiman, R. J., R. E. Bilby, D. E. Schindler, and J. M. Helfield. 2002. Pacific salmon, nutrients, and the dynamics of freshwater and riparian ecosystems. Ecosystems 5:399–417.

Nakagawa, S., and H. Schielzeth. 2013. A general and simple method for obtaining R2 from generalized linear mixed-effects models. Methods in Ecology and Evolution 4:133–142.

Novak, M. 2010. Estimating interaction strengths in nature: experimental support for an observational approach. Ecology 91:2394–2405.

Novak, M., C. Wolf, K. E. Coblentz, and I. D. Shepard. 2017. Quantifying predator dependence in the functional response of generalist predators. Ecology Letters 20:761–769.

Novak, M., and J. T. Wootton. 2008. Estimating nonlinear interaction strengths: an observation-based method for species-rich food webs. Ecology 89:2083–2089.

Paine, R. T. 1992. Food-web analysis through field measurement of per capita interaction strength. Nature 355:73–75.

Palmer, M. A., and N. L. Poff. 1997. The influence of environmental heterogeneity on patterns and processes in streams. Journal of the North American Benthological Society 16:169–173.

Peckarsky, B. L. 1983. Biotic interactions or abiotic limitations? A model of lotic community structure. Pages 303-323. Dynamics of Lotic Ecosystems. Ann Arbor Science. Ann Arbor MI.

Peckarsky, B. L., S. C. Horn, and B. Statzner. 1990. Stonefly predation along a hydraulic gradient: a field test of the harsh—benign hypothesis. Freshwater Biology 24:181–191.

Poisot, T., D. B. Stouffer, and D. Gravel. 2015. Beyond species: why ecological interaction networks vary through space and time. Oikos 124:243–251.

Polis, G. A. 1991. Complex trophic interactions in deserts: an empirical critique of food-web theory. The American Naturalist 138:123–155.

Polis, G. A., R. D. Holt, B. A. Menge, and K. O. Winemiller. 1996. Time, space, and life history: influences on food webs. Pages 435–460. Food webs. Springer. New York, NY.

Power, M. E., M. S. Parker, and W. E. Dietrich. 2008. Seasonal reassembly of a river food web: floods, droughts, and impacts of fish. Ecological Monographs 78:263–282.

Power, M. E., R. J. Stout, C. E. Cushing, P. P. Harper, F. R. Hauer, W. J. Matthews, P. B. Moyle, B. Statzner, R. Irene, and W. De Badgen. 1988. Biotic and abiotic controls in river and stream communities. Journal of the North American Benthological Society 7:456–479.

Preston, D. L., J. S. Henderson, L. P. Falke, and M. Novak. 2017. Using survival models to estimate invertebrate prey identification times in a generalist stream fish. Transactions of the American Fisheries Society 146:1303-1314.

Preston, D. L., J. S. Henderson, L. P. Falke, L. M. Segui, T. J. Layden, and M. Novak. 2018. What drives interaction strengths in complex food webs? A test with feeding rates of a generalist stream predator. Ecology 99:1591-1601.

Raffaelli, D. G., and S. J. Hall. 1996. Assessing the relative importance of trophic links in food webs. Pages 185–191 Food Webs. Springer. New York, NY.

Rall, B. C., U. Brose, M. Hartvig, G. Kalinkat, F. Schwarzmüller, O. Vucic-Pestic, and O. L. Petchey. 2012. Universal temperature and body-mass scaling of feeding rates. Phil. Trans. R. Soc. B 367:2923–2934.

de Ruiter, P. C., A.-M. Neutel, and J. C. Moore. 1995. Energetics, patterns of interaction strengths, and stability in real ecosystems. Science 269:1257.

Schleuning, M., N. Blüthgen, M. Flörchinger, J. Braun, H. M. Schaefer, and K. Böhning-Gaese. 2011. Specialization and interaction strength in a tropical plant–frugivore network differ among forest strata. Ecology 92:26–36.

Skalski, G. T., and J. F. Gilliam. 2001. Functional responses with predator interference: viable alternatives to the Holling type II model. Ecology 82:3083–3092.

Spiller, D. A., and T. W. Schoener. 2007. Alteration of island food-web dynamics following major disturbance by hurricanes. Ecology 88:37–41.

Tonkin, J. D., M. T. Bogan, N. Bonada, B. Rios-Touma, and D. A. Lytle. 2017. Seasonality and predictability shape temporal species diversity. Ecology 98:1201–1216.

Tylianakis, J. M., R. K. Didham, J. Bascompte, and D. A. Wardle. 2008. Global change and species interactions in terrestrial ecosystems. Ecology Letters 11:1351–1363.

Tylianakis, J. M., and R. J. Morris. 2017. Ecological networks across environmental gradients. Annual Review of Ecology, Evolution, and Systematics 48: 24-45.

Vander Zanden, M. J., J. M. Casselman, and J. B. Rasmussen. 1999. Stable isotope evidence for the food web consequences of species invasions in lakes. Nature 401:464-467.

Vázquez, D. P., S. B. Lomáscolo, M. B. Maldonado, N. P. Chacoff, J. Dorado, E. L. Stevani, and N. L. Vitale. 2012. The strength of plant–pollinator interactions. Ecology 93:719–725.

Warren, C. E., J. H. Wales, G. E. Davis, and P. Doudoroff. 1964. Trout production in an experimental stream enriched with sucrose. The Journal of Wildlife Management 28:617–660.

Wolf, C., M. Novak, and A. I. Gitelman. 2017. Bayesian characterization of uncertainty in species interaction strengths. Oecologia 184:327–339.

Woodward, G., J. P. Benstead, O. S. Beveridge, J. Blanchard, T. Brey, L. E. Brown, W. F. Cross, N. Friberg, T. C. Ings, U. Jacob, et al. 2010. Ecological networks in a changing climate. Advances in Ecological Research. 42:71-138.

Woodward, G., D. C. Speirs, A. G. Hildrew, and C. Hal. 2005. Quantification and resolution of a complex, size-structured food web. Advances in Ecological Research 36:85–135.

Wootton, J. T. 1997. Estimates and tests of per capita interaction strength: diet, abundance, and impact of intertidally foraging birds. Ecological Monographs 67:45–64.

Wootton, J. T., and M. Emmerson. 2005. Measurement of interaction strength in nature. Annual Review of Ecology, Evolution, and Systematics 36:419–444.

Wootton, J. T., M. S. Parker, and M. E. Power. 1996. Effects of disturbance on river food webs. Science 273:1558–1561.

Zuur, A. F, E. N. Ieno, N. Walker, A. A. Saveliev, G. M. Smith. 2009. Mixed effects models and extensions in ecology with R. Springer. New York, NY.

